# Significant Subgraph Mining for Neural Network Inference with Multiple Comparisons Correction

**DOI:** 10.1101/2021.11.03.467050

**Authors:** Aaron J. Gutknecht, Michael Wibral

## Abstract

We describe how the recently introduced method of significant subgraph mining can be employed as a useful tool in network comparison. It is applicable whenever the goal is to compare two sets of unweighted graphs and to determine differences in the processes that generate them. We provide an extension of the method to dependent graph generating processes as the occur for example in within-subject experimental designs. Furthermore, we present an extensive investigation of error-statistical properties of the method in simulation using Erdős-Rényi models and in empirical data. In particular, we perform an empirical power analysis for transfer entropy networks inferred from resting state MEG data comparing autism spectrum patients with neurotypical controls. From this analysis one may estimate that the appropriate sample size for similar studies should be chosen in the order of n=60 per group or larger.

**Author summary:** A key objective of neuroscientifc research is to determine how different parts of the brain are connected. The end result of such investigations is always a graph consisting of nodes corresponding to brain regions or nerve cells and edges between the nodes indicating if they are connected or not. The connections may be structural (an actual anatomical connection) but can also be functional – meaning that there is a statistical dependency between the activity in one part of the brain and the activity in another. A prime example of the latter type of connection would be the information flow between brain areas. Differences in the patterns of connectivity are likely to be responsible for and indicative of various neurological disorders such as autism spectrum disorders. It is therefore important that efficient methods to detect such differences are available. The key problem in developing methods for comparing patterns of connectivity is that there is generally a vast number of different patterns (it can easily exceed the number of stars in the milky way). In this paper we describe how the recently developed method of significant subgraph mining accounts for this problem and how it can be usefully employed in neuroscientifc research.

## Introduction

Comparing networks observed under two or more different conditions is a pervasive problem in network science in general, and especially in neuroscience. The basic question in all of these cases is if the observed patterns or motifs in two samples of networks differ solely due to chance or because of a genuine difference between the conditions under investigation. For example, a researcher may ask if a certain pattern of functional connections in a brain network reconstructed from magnetoencephalography (MEG) data is more likely to occur in individuals with autism spectrum disorder than in neurotypic controls, or whether an observed difference in occurrence is solely due to chance. What makes this question difficult to answer is the fact that the number of possible patterns in the network scales as 2^*l*^2^^, with *l* the number of network nodes, – leading to a formidable multiple comparison problem. Correcting for multiple comparisons with standard methods (e.g. Bonferroni) typically leads to an enormous loss of power as these methods do not exploit the particular properties of the network comparison problem. We here show how to utilize a recently developed mathematical approach, called Significant Subgraph Mining (SSM) [1,2], to efficiently solve the network-comparison problem while maintaining strict bounds on type *I* error rates for between unit of observation designs. We then also extend this method to the case of within unit of observation designs, that was, to our best knowledge, not described in the literature before. In this study we will explain the core idea behind these methods for a broad audience, but also provide a rigorous mathematical exposition for reference. We furthermore provide a detailed study of the error-statistical properties (family-wise error rate and statistical power) in both simulation and empirical data.

## Background and Theory

Neural networks can usefully be described as graphs consisting of a set of nodes and a set of edges connecting the nodes. The nodes represent specific parts of the network such as individual neurons, clusters of neurons, or larger brain regions, whereas the edges represent relationships between these parts. Depending on whether the relationship of interest is symmetric (such as correlation) or asymmetric (such as Transfer Entropy or Granger Causality) the network can be modelled as an undirected or as a directed graph respectively. Once we have a decided upon an appropriate graph theoretic description, we can apply it to neural networks measured in two different experimental groups resulting in two sets of graphs. In doing so we are essentially sampling from two independent *graph-generating processes* (see Figure 1 for illustration).

**Fig 1.**
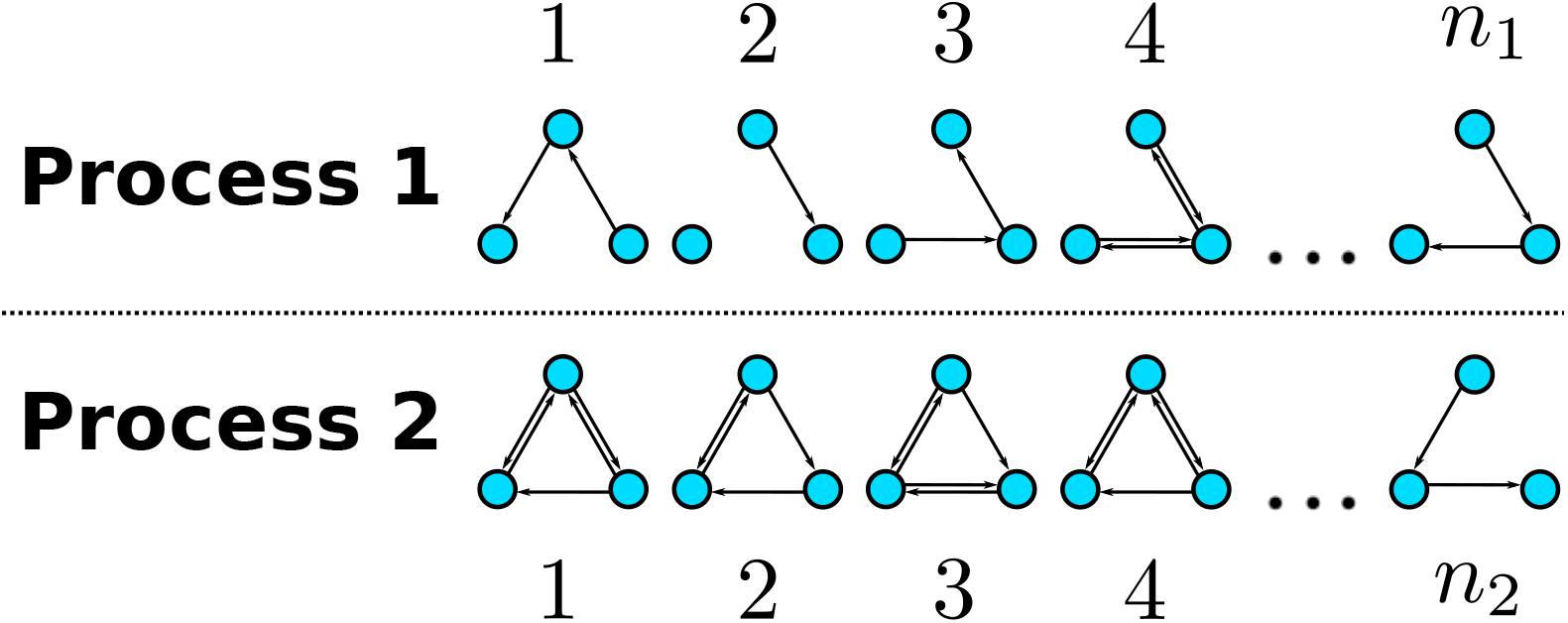
Illustration of two graph-generating processes. Each process consists of randomly sampling individuals from a specific population and describing the neural activity of these individuals as a graph. The population underlying process 1 is sampled *n*_1_ times and the population underlying process 2 is sampled *n*_2_ times. The nodes may correspond to different brain areas while the edges describe any directed relationship between brain areas such as information transfer.

The key question is now if there are any significant differences between these two sets. However, since graphs are complex objects it is not immediately obvious how they should be compared. In principle one may imagine numerous different possibilities, for instance, comparing the average number of connections that a node has or the average number of steps it takes to get from one node to another. Instead of relying on such summary statistics, however, one may also take a more fine-grained approach by looking for differences with respect to any possible pattern, or more technically *subgraph,* that may have been observed. Does a particular edge occur significantly more often in one group than in the other? What about particular bi-directional connections? Or are there even more complex subgraphs -consisting of many links-that are more frequently observed in one of the groups? Answering such questions affords a particularly detailed description of the differences between the two processes. In particular, it not only allows us to *detect* such differences but also to *localize* them. If a significant difference with respect to a particular subgraph is found, then we have reason to believe that there is a difference between the probabilities with which this subgraph occurs in the two groups.

Figure 2 shows examples of different subgraphs of a graph with three edges.

**Fig 2.**
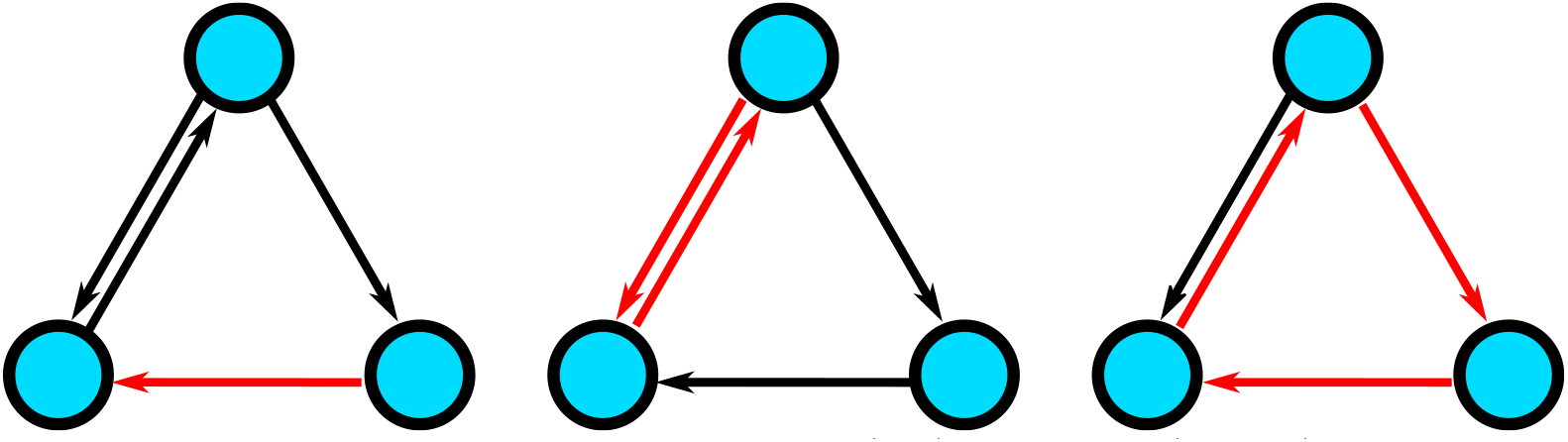
Illustration of subgraphs with one edge (left), two edges (middle), and three edges (right) of a graph with three nodes.

The process of enumerating all subgraphs for which there is a significant difference between the groups is called *significant subgraph mining* [1]. The goal is to identify all subgraphs that are generated with different probabilities by the two processes. The main difficulty underlying significant subgraph mining is that the number of possible subgraphs grows extremely quickly with the number of nodes. For a directed graph with seven nodes, it is already in the order of 10^14^. This not only imposes runtime constraints but also leads to a severe multiple comparisons problem. Performing a significance test for each potential subgraph and then adjusting by the number of tests is not a viable option because the resulting test will have an extremely poor statistical power. As will be detailed later, due to the discreteness of the problem the power may even be exactly zero because p-values low enough to reach significance can in principle not be achieved. In the following sections we will formally introduce significant subgraph mining by first setting up an appropriate probabilistic model, explaining how to construct a significance test for a particular subgraph, and finally, detailing two methods for solving the multiple comparisons problem.

### Probabilistic Model

We are considering two independently sampled sets of directed graphs 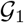 and 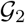 describing, for instance, connections between brain regions in two experimental groups. Each set contains one graph per subject in each group and we assume that the (fixed) sample sizes of each group are 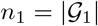 and 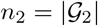. All graphs are defined on the same set of nodes *V* = {1, 2,..., *l*} but may include different sets of links (edges) *E* ⊆ *V* × *V*. The graphs are assumed to have been generated by two potentially different graph-generating processes. Each process can be described by a random *l × l* adjacency matrix of, possibly dependent, Bernoulli random variables. Each of those variables tells us whether a particular link is present (“1”) or absent (“0”) in the generated graph:

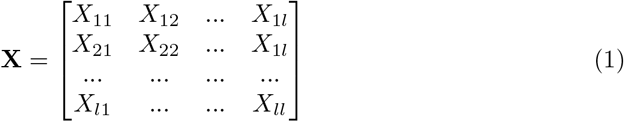

where

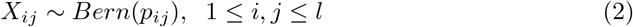

A graph-generating process can be fully characterized by the probabilities with which it generates possible subgraphs. Specifically, there is one such probability for each subgraph of the fully connected graph *G_C_* = (*V, V × V*): If *G* = (*V_G_, E_G_*) ⊆ *G_C_*, then the probability that G occurs as a subgraph of the generated graph is given by

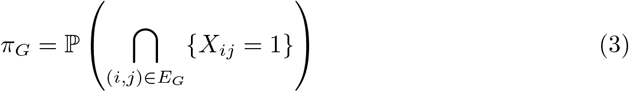

where (*i,j*) indicates an individual link from node *i* to node *j*. It is important to note that *π_G_* denotes the the probability that all the edges of G are realized *plus possibly some additional edges.* This is to be distinguished from the probability that *exactly* the graph *G* is realized. In the following we will always refer to the probability *π_G_* as the *subgraph probability* of *G*. The sequence of subgraph probabilities *π* = (*π_G_*)_*G*⊆*G_c_*_ provides a complete probabilistic description of a graph generating process. We can now model the two sets of directed graphs 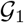 and 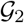 as realizations of two independent graph generating processes **X**^(1)^ and **X**^(2)^:

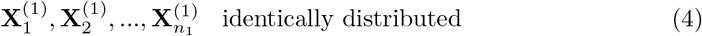

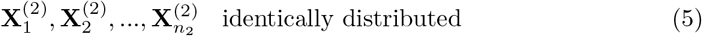

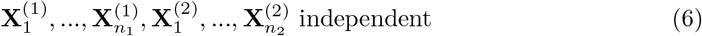

where each variable is a random *l × l* adjacency matrix representing a particular graph. Process **X**^(1)^ generates graphs according to subgraph probabilities *π*^(1)^ whereas the subgraph probabilities for process **X**^(2)^ are given by *π*^(2)^. Based on this probabilistic model we may now proceed to test for differences between the two processes.

### Testing Individual Subgraphs

Our goal is now to find those subgraphs G that are generated with probabilities that are different for the two processes. If the two processes describe two distinct experimental groups, this means that we are trying to identify subgraphs whose occurrence depends on group membership. Thus, for each possible subgraph G, we are testing the null hypothesis of *equal subgraph probabilities,* or equivalently, of *independence of subgraph occurrence from group membership*

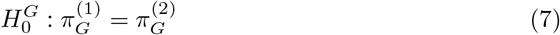

against the alternative of unequal subgraph probabilities or dependence on group membership

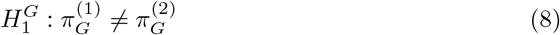

In order to test such a null-hypothesis we have to compare how often the subgraph G occurred in each group and determine if the observed difference could have occurred by chance, i.e. if the probability of such a difference would be larger than *α* under the null-hypothesis. The relevant data for this test can be summarized in a 2 × 2 contingency table:

| Subgraph G | Occurrences | Non-Occurrences | Total |
| --- | --- | --- | --- |
| Group 1 | $f_1(G)$ | $n_1 - f_1(G)$ | $n_1$ |
| Group 2 | $f_2(G)$ | $n_2 - f_2(G)$ | $n_2$ |
| Total | $f(G)$ | $n - f(G)$ | $n$ |

where *f_i_*(*G*) denotes the *observed* absolute frequency of subgraph G in Group i, *f* (*G*) = *f*_1_(*G*) + *f*_2_(*G*) denotes the *observed* absolute frequency of G in the entire data set, and *n = n*_1_ + *n*_2_ is the total sample size. In the following, we will use *F_i_*(*G*) and *F*(*G*) to denote the corresponding *random* absolute frequencies. Given our model assumptions above, the numbers of occurrences in each group are independent Binomial variables: On each of the *n*_1_ (or *n*_2_) independent trials there is a fixed probability 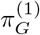 (or 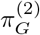) that the subgraph G occurs. Thus,

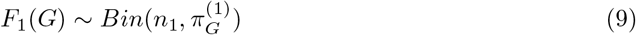

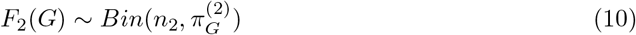

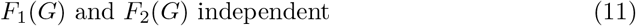

This means that our goal is to compare two independent Binomial proportions. In principle, there is a large number of possible tests of the null-hypothesis of interest [3,4]. In the following, we will utilize Fisher’s exact test as has also been done in previous studies [1,2]. This test has the advantage that it does not require any minimum number of observations per cell in the contingency table. By contrast, well known asymptotic tests such as Pearson’s χ^2^-test are only valid in large samples. This is problematic in a subgraph mining context because in practice many subgraphs will only occur very rarely, or sometimes not at all.

The key idea underlying Fisher’s exact test is to condition on the total number of occurrences *f* (*G*): Given that the subgraph G occurred *f* (*G*) times in the entire data set, we would expect these occurrences to be distributed across the two groups proportional to the respective sample sizes if the subgraph probability of G is actually equal in both groups. For instance, if the sample sizes are equal, then we would expect roughly half of the occurrences to be observed in group 1 and roughly half of the occurrences to be observed in group 2. This means that, given the total number of occurrences *f* (*G*), the number of occurrences in the fist group *f*_1_(*G*) serves as a reasonable test statistic. Specifically, the random variable *F*_1_(*G*) can be shown to follow a hypergeometric distribution under the null-hypothesis and conditional on the total number of occurrences:

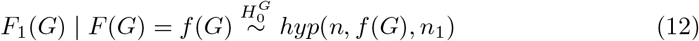

In other words, if the null-hypothesis is true and given the total number of occurrences, the *n*_1_ occurrences and non-occurrences of subgraph G in Group 1 are assigned as if they were drawn randomly without replacement out of an urn containing exactly *f* (*G*) occurrences and *n* – *f* (*G*) non-occurrences (see Figure 3). *F*_1_(*G*) can now be used as a test-statistic for the hypothesis test. Since we are interested in differences between the graph generating processes in either direction the appropriate test is a *two-sided* one. One way to construct such a test is two combine two one-sided tests in the following way: For a right-sided test of the null-hypothesis against the alternative 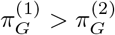 the p-value can be computed as

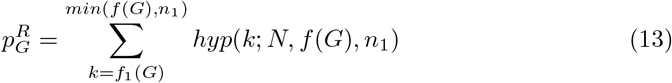

summing up the probabilities of all possible values of *f*_1_(*G*) *larger than or equal to* the one actually observed. Note that *f*_1_(*G*) cannot be larger than *min*(*f* (*G*),*n*_1_) because the number of occurrences in Group 1 can neither be larger than the sample size *n*_1_ nor larger than the total number of occurrences *f* (*G*). Similarly, the left-sided p-value against the alternative 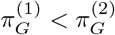 is

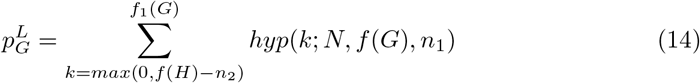

summing up the probabilities of all possible values of *f*_1_(*G*) *smaller than or equal to* the one actually observed. Here the summation starts at *max*(0, *f* (*H*) – *n*_2_) because, first, the number of occurrences cannot be negative, and second, if the total number of occurrences is larger than the sample size of the second group, then there must be at least *f* (*H*) – *n*_2_ occurrences in the first group.

**Fig 3.**
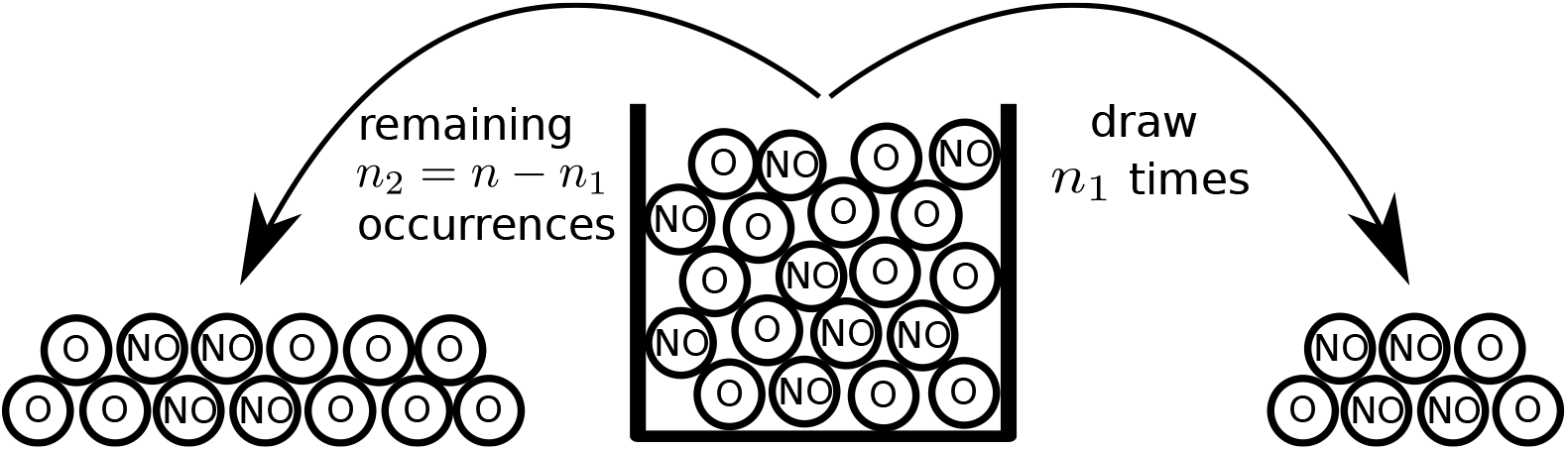
Comparing two Binomial proportions using Fisher’s exact test. Under the null-hypothesis and conditional on the total number of occurrences of a subgraph, the occurrences are distributed over the groups as if drawn at random without replacement out of an urn containing one ball per subject. The balls are labelled ‘O’ if the subgraph occurred in the corresponding subject and ‘NO’ if it did not occur. In the illustration *n* = 20 (number of total measurements, balls), *n*_1_ = 7 (number of measurements for group 1, black balls), and *f* (*G*) = 12 (number of occurences, balls with ‘O’). The seven balls drawn for group 1 are shown to the right of the urn. They include three occurrences and four non-occurrences. This result would lead to an insignificant p-value of ≈ 0.5

The idea is now to reject the null-hypothesis against the two-sided alternative just in case at least one of the one-sided tests turns out to be significant. Keeping in mind that we have to correct for having performed two tests by doubling the one-sided p-values,this means that the null-hypothesis should be rejected if and only if the two-sided p-value

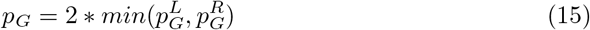

is smaller than or equal to *α*^1^. In order to make sure that the resulting test is valid it is worthwhile to check the probability of rejection under the null-hypothesis:

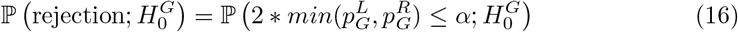

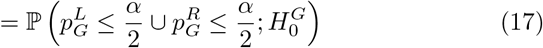

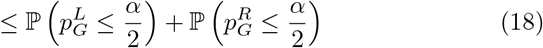

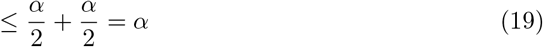

### Multiple Comparisons

Since there may be a very large number of possible subgraphs to be tested we are faced with a difficult multiple comparisons problem. For a directed graph with 7 nodes the number of possible subgraphs is already in the order of 10^14^. If we were to use this number as a Bonferroni correction factor the testing procedure would have an exceedingly low statistical power meaning that it would be almost impossible to detect existing differences in subgraph probabilities. In the following, we will describe two methods for solving the multiple comparisons problem: the Tarone correction [5] and the Westfall-Young permutation procedure [6]. Both methods have previously been applied in a subgraph mining context [1, 2]. The two methods are *single-step methods* meaning that they prescribe a single corrected significance threshold *δ* against which the p-values of all hypotheses are compared [6]. A hypothesis is rejected just in case its p-value falls below this threshold. We will now consider the Tarone and Westfall-Young corrections in turn.

#### Tarone’s Correction

The subgraph mining problem is discrete in the sense that there is only a finite number of possible p-values. This fact can be exploited to drastically reduce the correction factor. The key insight underlying the Tarone correction is that for any total frequency *f* (*G*) of a particular subgraph *G* there is a *minimum achievable p-value* given *F*(*G*) = *f* (*G*) which we will denote by 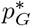. Intuitively, this minimum achievable p-value is reached if the *f* (*G*) occurrences are distributed as unevenly as possible between the two groups. We may now introduce the notion of the set *T*(*k*) of 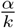-testable subgraphs:

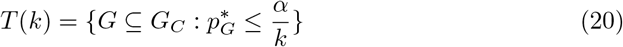

containing all subgraphs whose minimum achievable p-value is smaller than or equal to 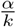. Following Tarone, the number of elements of this set can be denoted by *m*(*k*) = |*T*(*k*)|. Tarone et al then showed that the smallest integer *k* such that 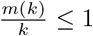 is a valid correction factor and that this number is often considerably smaller than the Bonferroni-factor [5]. We will call the Tarone correction factor *K*(*α*) in the following. Figure 4 illustrates the concepts of testable, untestable, and significant subgraphs.

**Fig 4.**
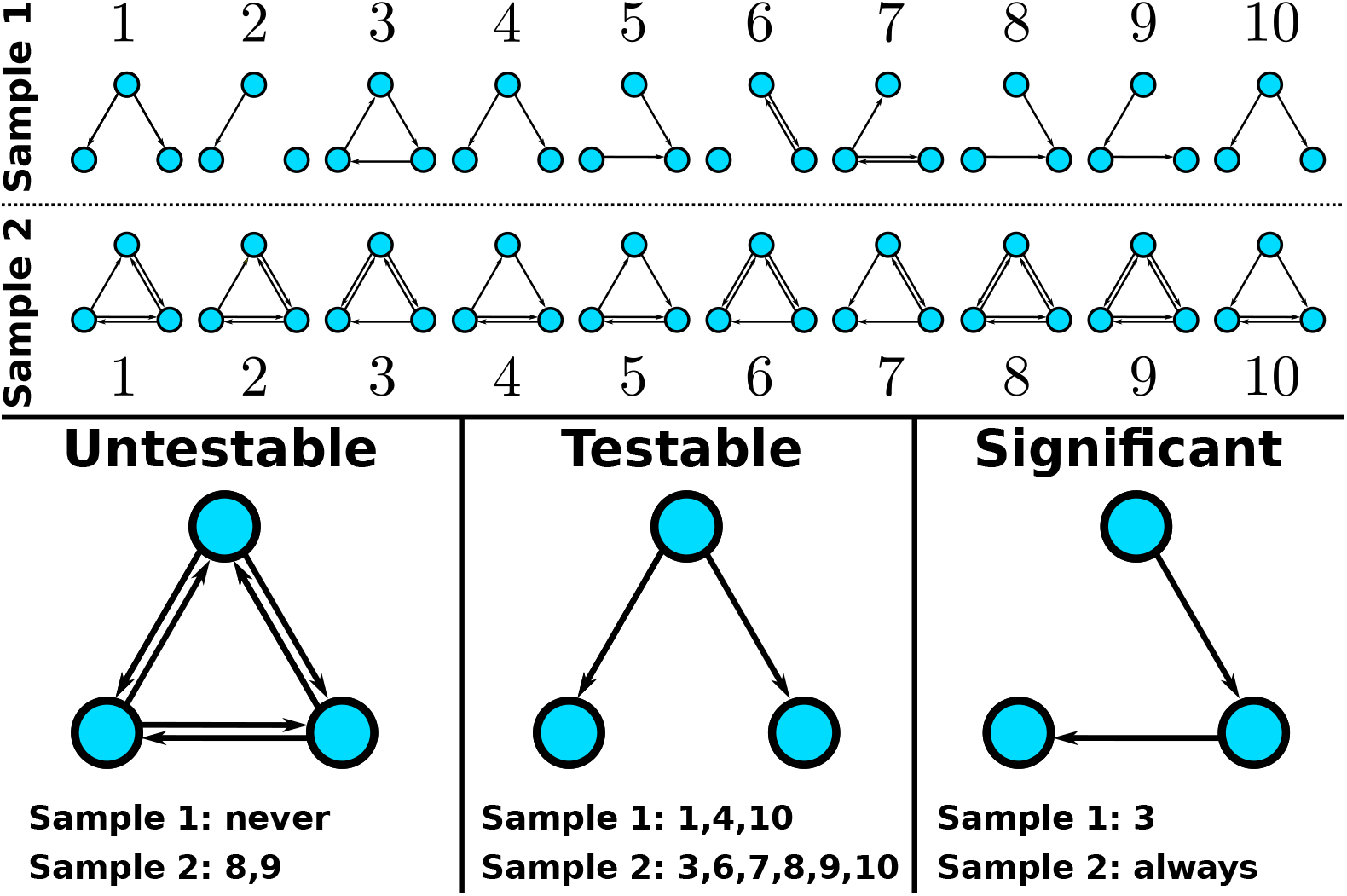
Examples of 0.05-untestable, 0.05-testable, and significant subgraphs for a data set consisting of 10 graphs per group (top panel). The fully connected graph is untestable at level 0.05 because it only occurs twice in the data set (group 2 samples 8 and 9) leading to a minimum achievable p-value of ≈ 0.47. The graph shown on the bottom middle is testable at level 0.05 since it occurs 9 times in total. This means that its minimum achievable p-value is ≈ 0. 0001. However, it is not significant with an actual (uncorrected) p-value of ≈ 0.37. The graph shown on the bottom right reaches significance using Tarone’s correction factor *K*(0.05) = 17. It occurs every time in group 2 but only once it group 1 which results in a corrected p-value of ≈ 0.02.

The validity of *K*(*α*) can be seen as follows: Let 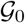 denote the set of subgraphs for which the null hypothesis of equal subgraph probabilities is true and let 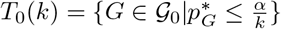 be the subset of 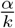-testable subgraphs within 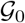. Furthermore, let *m*_0_(*k*) be the number of elements of this set, i.e. the number of 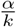-testable subgraphs for which the null-hypothesis is true. We can now compute the conditional family-wise error rate for a correction factor *k* ∈ ℕ given the observed total frequencies of each subgraph. These frequencies can be interpreted as the realization of a random vector F containing one entry *F*(*G*) per possible subgraph:

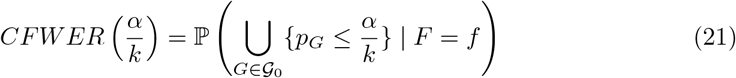

We only have to take the union over 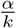-testable subgraphs because all other terms have probability zero:

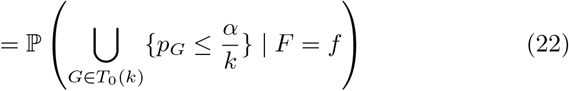

Using Boole’s inequality (the “union bound”):

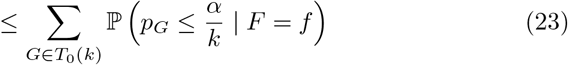

By construction of the p-value we have for any constant 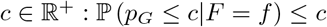 (include proof in appendix?). This fact can be applied to each term in the above sum with 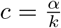:

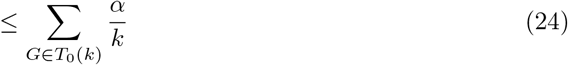

The sum has *m*_0_(*k*) terms:

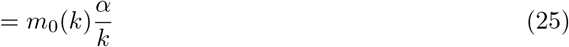

The number of testable subgraphs for which the null-hypothesis is true is smaller than or equal to the total number of testable subgraphs:

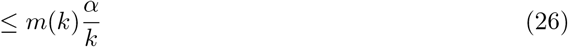

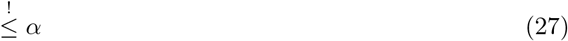

For the final equation (6) to be valid it must be true that 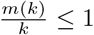. In order to maximize the power of the resulting test, the correction factor should be chosen as small as possible. Hence, the appropriate choice is the smallest integer k such that 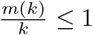, i.e. *K*(*α*). Since the argument is valid for all possible observed total frequencies, it is also valid for the unconditional FWER which is simply a weighted average of the conditional FWERs:

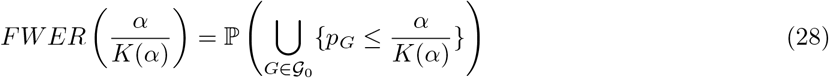

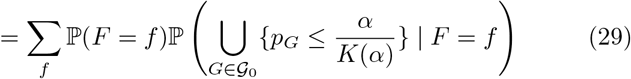

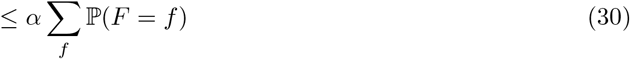

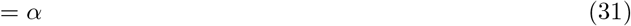

It is important to note that this argument does not make any assumptions about *which* or *how many* null-hypotheses are in fact true. The FWER is controlled in all cases. This property is called *strong control* of the FWER.

The Tarone correction has been criticized on the basis that it is not *α*-consistent [7]. This means that a null-hypothesis might not be rejected at level *α* even though it would have been rejected at an even smaller level *δ < α*. However, there is a simple modification proposed by Hommel et al [8] that makes the Tarone procedure *α*-consistent and, maybe more importantly, also improves its statistical power. The idea is to make the procedure *α*-consistent *by definition,* i.e. to reject 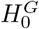 if the standard Tarone procedure would reject or if there exists a level *γ < α* such that the standard Tarone procedure would reject:

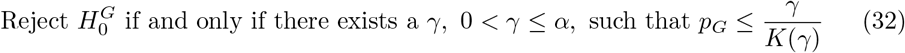

This rule has to be at least as powerful as the standard Tarone procedure because a null-hypothesis is rejected by the standard procedure it is also rejected by the improved version. Additionally, there are cases in which the Hommel impovement rejects but the standard Tarone procedure does not. Hommel presented a simple algorithm to implement this idea which, in the subgraph mining context, can be phrased as follows: First, we order all subgraphs in terms of their minimal achievable p-values such that 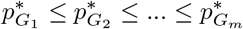, where *m* = 2^*l*^2^^ is the total number of possible subgraphs. Then we define the rejection rule as:

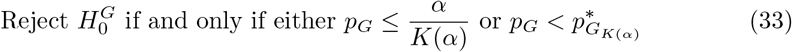

In the next section we will explain a second possible multiple comparisons correction which is based on a permutation strategy: the Westfall-Young correction [2, 6].

#### Westfall-Young Correction

The familiy-wise error rate with respect to a significance level *δ* can be expressed in terms of the CDF of the smallest p-value associated with a true null-hypothesis: The event that there is at least one false positive is identical with the event that the smallest p-value associated with a true null-hypothesis is smaller than *δ*. Accordingly we have for the conditional FWER:

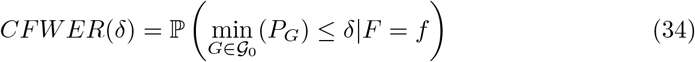

This means that if the correction factor is chosen as the *α*-quantile of the conditional distribution of 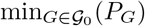 given the observed total occurrences, then the CFWER is bounded by α. The problem is that we cannot evaluate the required distribution because we don’t know which hypotheses are true. The idea underlying the Westfall-Young correction is to instead define the correction factor as the *α*-quantile of the distribution of the minimal p-value across *all* subgraphs and under the *complete* null-hypothesis:

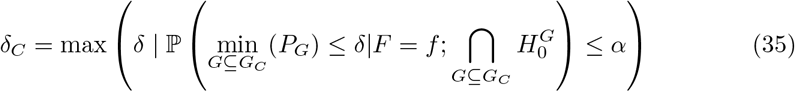

where the maximum is taken over all possible p-values. This correction factor always provides *weak control* of the FWER in the sense that the FWER is bounded by *α* under the complete null-hypothesis (we will address the issue of strong control below). In this scenario we have that 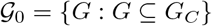 and hence

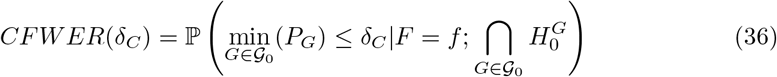

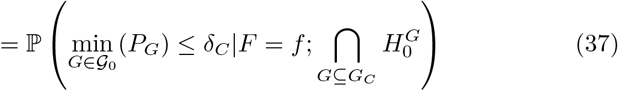

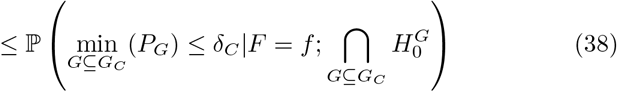

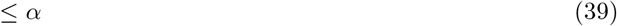

The quantile *δ_C_* and its underlying distribution can now be estimated via permutation strategies. The procedure is as follows: First, we may represent the entire data set by the following table

| Subject | Group | $G_1$ | $G_2$ | ... | $G_m$ |
| --- | --- | --- | --- | --- | --- |
| 1 | 0 | 0 | 1 | ... | 1 |
| 2 | 0 | 1 | 1 | ... | 1 |
| ... | ... | ... | ... | ... | ... |
| $n_1$ | 0 | 0 | 0 | ... | 1 |
| $n_1 + 1$ | 1 | 1 | 1 | ... | 1 |
| ... | ... | ... | ... | ... | ... |
| $n_1 + n_2$ | 1 | 0 | 1 | ... | 1 |

The columns labelled *G_i_* tell us if subgraph *G_i_* was present or absent in the different subjects (rows). The column labelled “Group” describes which group the different subjects belong to. Under the complete null-hypothesis the group labels are arbitrarily exchangeable. This is because, given our independence assumptions (see Equation 6), all the observed graphs in the data set are independent and identically distributed samples from the same underlying distribution in the complete null-case:

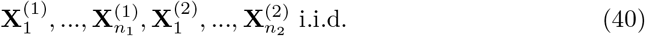

Accordingly, the distribution of these random matrices is exchangeable. The column of groub labels is now shuffled, reassigning the graphs in the data set to the two groups. Based on this permuted data set we can recompute a p-value for each *G_i_* and determine the smallest of those p-values. Repeating this process many times allows us to obtain a good estimate of the distribution of the smallest p-value under the complete null-hypothesis. The resulting distribution is the desired distribution in equation 34, because given F=f everything is fixed except the allocation of group labels (and the allocation of subject labels but those do not affect the p-values). Thus, the conditional distribution of the minimal p-value under the complete null-hypotheses and given F=f is identical to the permutation distribution:

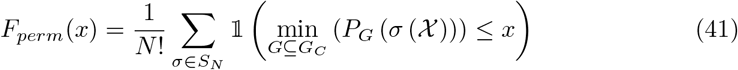

where *S_N_* is the group of permutations on N elements, 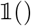 is the indicator function, and 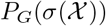 is the p-value associated with subgraph G *with respect to the permuteted data set* 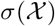. In other words, the permutation distribution evaluated at a point x simply measures the fraction of permutations σ for which the smallest p-value across all subgraphs is smaller than or equal to x. Since the number of permutations grows very quickly with the total sample size N it is usually not possible to evaluate all permutations. Instead, one has to consider a much smaller random sample of permutations in order to obtain an approximation to the permutation distribution. This procedure can be shown to be valid as long as the identity permutation (i.e. the original data set) is always included [9].

Now, ideally we would like the FWER to be controlled no matter how many and which null-hypotheses are true, i.e. we wish to obtain *strong control* of the FWER. In order to achieve strong control one would generally have to consider all possible ‘intersection hypotheses’, i.e. logical conjunctions of null-hypotheses [10]. However, since any hypothesis can be either included or excluded in the conjunction, there are 2^*m*^ possible conjunctions, where m is the number of hypotheses. Since for a simple directed graph with l nodes *m* = 2^*l*^2^^, this number grows extremely quickly. Even for only three nodes it is already in the order of 10^154^, and hence, the problem becomes computationally intractable. In order to achieve strong control without having to consider all intersection hypotheses an additional assumption called *subset pivotality* can be utilized [6,11]. Formally, a vector of p-values **P** = (*P*_1_,*P*_2_, …,*P_m_*) has subset pivotality if and only if for any subset of indices *K* ⊆ {1, 2,..., *m*} the joint distribution of the subvector *P_K_* = {*P_i_*|*i* ∈ *K*} is the same under the restrictions 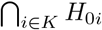 and 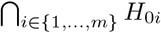 [6]. In other words, the joint distribution has to be same in both of the following scenarios:

1. All null-hypotheses associated with the p-values in the subvector are true.
2. All null-hypotheses are true.

Appropriate conditional versions of subset pivotality given F = f can be defined analogously by demanding the same property to be true for the conditional joint distribution. In the subgraph mining context subset pivotality means in particular that the the joint distribution of p-values corresponding to subgraphs for which the null-hypothesis is in fact true remains unchanged in the (possibly counterfactual) scenario that the null-hypothesis is true for *all* subgraphs. This implies that the distribution of the smallest p-value remains unchanged as well. If the underlying probabilistic model satisfies subset pivotality the above argument for the boundedness of the FWER (equations 36–39) remains valid even if only a subset of null-hypotheses are true. To our knowledge it is an open question whether subset pivotality may be satisfied in the subgraph mining context under suitable conditions (perhaps after imposing additional constraints on the graph-generating mechanisms). However, we suspect that it is not satisfied in general.

### Extension to Within-Subject Designs

So far we have considered networks associated with subjects from two groups and we assumed that the numbers of occurrences of a subgraph in the two groups are independent of each other. However, there are many cases in which there is only a single group of subjects and we are interested in how the networks differ between two experimental conditions. Since the same subjects are measured in both conditions, the independence assumption is not warranted anymore. Because Fisher’s exact test assumes independence, the approach described above has to be modified. In particular, in case of dependence, the null-distribution of the number of occurrences in the first group / condition will in general not be a hypergeometric distribution potentially leading to inflated type I error rates in Fisher’s exact test. An appropriate alternative is McNemars test for marginal homogeneity. It essentially tests the same null-hypothesis as Fisher’s exact test, but is based on a wider probabilistic model of the graph generating processes. In particular, the independence assumption is relaxed allowing for dependencies between the two experimental conditions: Whether a subgraph occurs in condition A in a particular subject may affect the probability of its occurrence in condition B and vice versa. Suppose we are observing n subjects in two conditions. We may denote the random adjacency matrices corresponding the i-th subject in condition 1 and 2 by 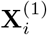 and 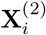, respectively. Then the probabilistic model for the graph-generating processes is:

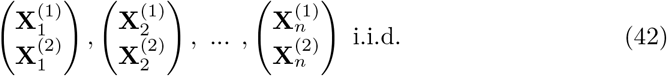

For each subject there is an independent and identically distributed realization of the two graph-generating processes. The two processes themselves may be dependent since they describe the same subject being observed under two conditions. The distributions of 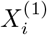 and 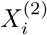 are again determined by the sequences of subgraph probabilities *π*^(1)^ and *π*^(2)^. Again, for any particular G we would like to test the null-hypothesis:

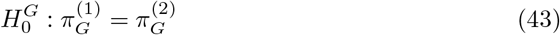

The idea underlying McNemar’s test is to divide the possible outcomes for each subject into four different categories: 1) G occurred in both conditions, 2) G occurred in neither condition, 3) G occurred in condition 1 but not in condition 2, 4) G occurred in condition 2 but not in condition 1. The first two categories are called *concordant pairs* and the latter two are called *discordant pairs*. The discordant pairs are of particular interest because differences in subgraph probabilities between the two conditions will manifest themselves in the relative number of the two types of discordant pairs: If 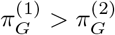 then we would expect to observe the outcome ‘G occurred only in condition 1’ more frequently than the outcome ‘G occurred only in condition 2’. Conversely, if 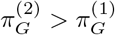, than we would expect to observe the latter type of discordant pair more frequently. The frequency of any of the four categories can be represented in a contingency table:

| Condition 1 / Condition 2 | Yes | No | Total |
| --- | --- | --- | --- |
| Yes | $Y_{11}^G$ | $Y_{10}^G$ | $F_1(G)$ |
| No | $Y_{01}^G$ | $Y_{00}^G$ | $n - F_1(G)$ |
| Total | $F_2(G)$ | $n - F_2(G)$ | $n$ |

The variables 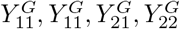 are the counts of the four categories. The numbers of occurrences in each condition *F*_1_(*G*) and *F*_2_(*G*) appear in the margins of the contingency table. McNemar’s test uses the upper right entry, 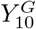, as the test-statistic. Conditional on the total number of discordant pairs, 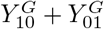, and under the null-hypothesis, this test-statistic has a binomial distribution

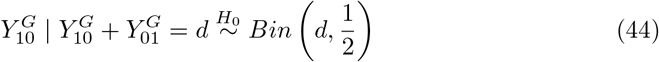

If there are exactly d discordant pairs and the probability of G is equal in both conditions, then both types of discordant pairs (‘only in condition 1’ or ‘only in condition 2’) occur independently with equal probabilities in each of the d subjects where a discordant pair was observed. A two-sided test can be constructed in just the same way as described above for the between-subjects case. First, we construct right- and left-sided p-values as:

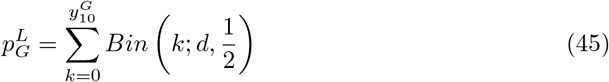

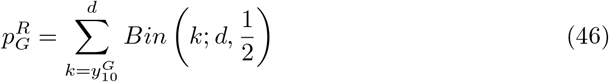

Then the two-sided p-value is

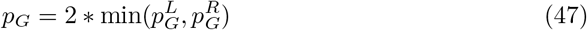

Exactly like the Fisher’s test, McNemar’s test also has a minimal achievable p-value. The only difference is that it is not a function of the total number of occurrences in condition A, but a function of the number of discordant pairs. The Tarone and Hommel corrections described above remain valid if Fisher’s exact test is simply replaced by McNemar’s test. The Westfall-Young procedure requires some modifications because the permutation strategy described above is not valid anymore. The problem is that, because of possible dependencies between the conditions, condition labels are not arbitrarily exchangeable under the complete null-hypothesis. Instead we have to take a more restricted approach and only exchange condition labels *within subjects*. In doing so we are not only keeping the total number of occurrences *F*(*G*) constant for each subgraph, but also the total number of discordant pairs *D*(*G*). Accordingly, the theoretical Westfall-Young correction factor, estimated by the modified permutation strategy, is the *α*-quantile, *δ_C_*, of the conditional distribution of the smallest p-value given *F = f* and *D* = *d* and under the complete null-hypothesis. The validity of *δ_C_* can be seen in just the same way as described above:

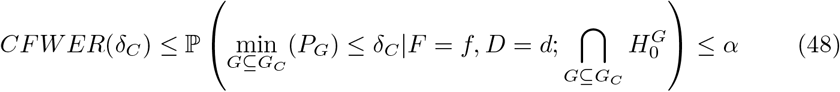

### Validation of Multiple Comparisons Correction Methods using Erdos-Renyi Models

In this section we empirically investigate the family-wise error rate and statistical power of the multiple comparison correction methods for significant subgraph mining described above. In doing so we will utilize Erdős-Rényi models for generating random graphs. In these models the edges occurs independently with some common probability p. in each graph-generating process. This means that the subgraph probability for a graph *G* = (*V_G_*, *E_G_*) in process i is *p_i_* raised to the number of edges G consists of:

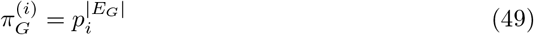

If *p_i_* is the same for both graph-generating processes (*p*_1_ = *p*_2_), then the complete null-hypothesis is satisfied. By contrast, if p is chosen differently for the two processes (*p*_1_ ≠ *P*_2_), then the null-hypothesis of equal subgraph probabilities is violated for all subgraphs, i.e. the *complete alternative* is satisfied. We used the former setting for the purposes of FWER estimation and the latter for power analysis. Furthermore, the two graph-generating processes were simulated independently of each other which corresponds to the between-subjects case. Accordingly, Fisher’s exact test was used throughout.

#### Family-Wise Error Rate

In order to empirically ascertain that the desired bound on the family-wise error rate are maintained by the Tarone and Westfall-Young correction in the subgraph mining context we performed a simulation study based on Erdos-Renyi models. We tested sample sizes *n* = 20, 30, 40, network sizes *l* = 2, 4, 6, 8, 10, and connection densities p = 0.1, 0.2, 0.3. For each combination of these values we carried out 1000 simulations and estimated the empirical FWER as the proportion of simulations in which one or more significant subgraphs were identified. Figure 5 shows the results of this analysis. The FWER is below the prespecified α = 0.05 in all cases. The Bonferroni correction is most conservative. In fact, the FWER quickly drops to exactly zero since the Bonferroni-corrected level is smaller than the smallest possible p-values. The Tarone-correction reaches intermediate values of 0.1-0.3 while the Westfall-Young correction is always closest the prespecified level and sometimes even reaches it.

**Fig 5.**
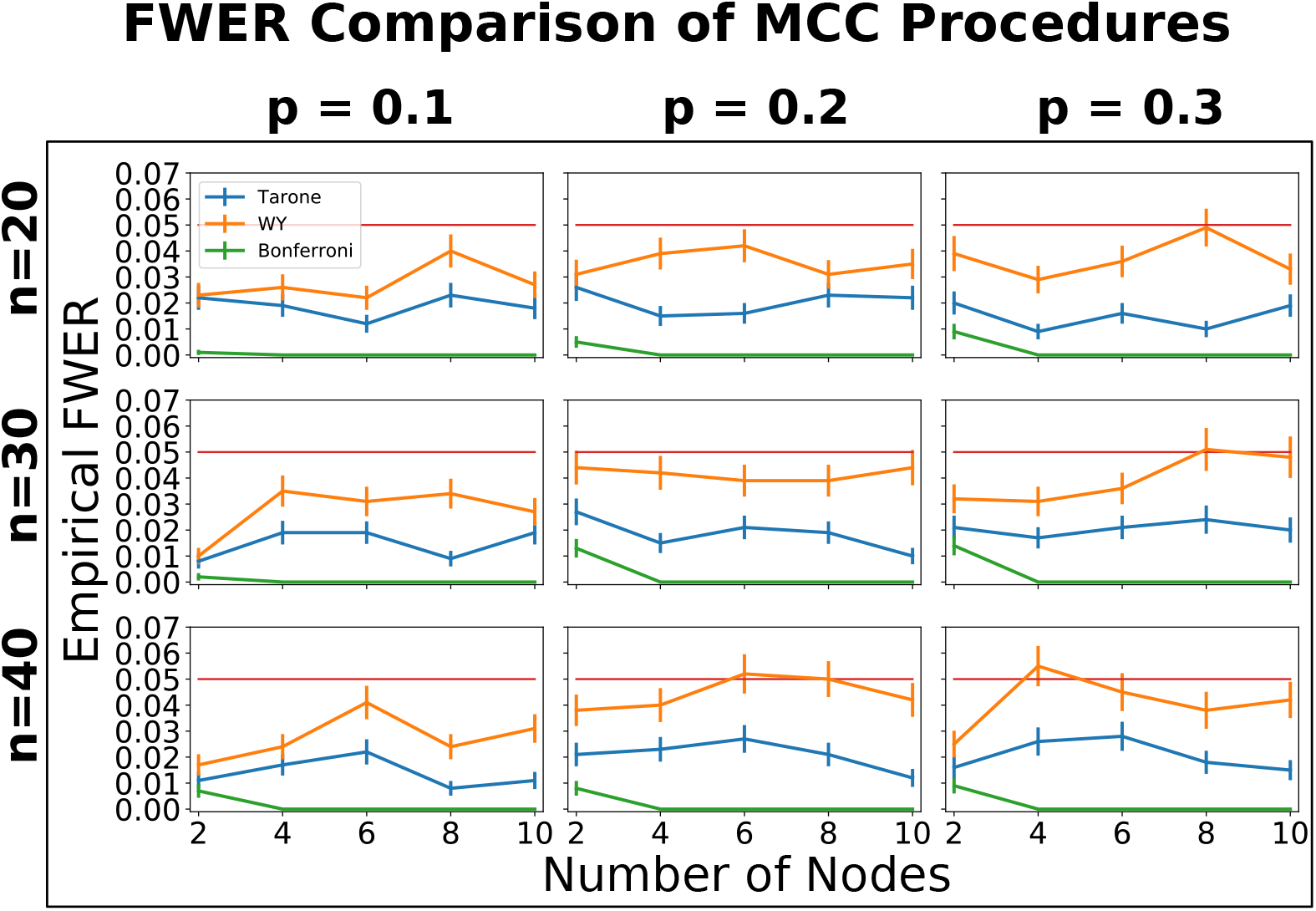
Estimated family-wise error rates of Tarone, Westfall-Young, and Bonferroni corrections based on 1000 simulations and different sample sizes, connection densities, and network sizes. Error-bars represent one standard-error. The estimated FWER never exceeded the desired FWER of *α* = 0.05 (red horizontal line) by more than one standard-error for all correction methods. In fact, it was always smaller than 0.05 except in three cases for the Westfall-Young correction (0.051, 0.052, and 0.055). The estimated FWERs of the three methods were always ordered in the same way: The Bonferroni correction had the smallest estimated FWER (at most 0.014), followed by the Tarone correction (at most 0.028), and the Westfall-Young correction (at most 0.055).

#### Power

We now turn our attention to the statistical power of the multiple comparison correction methods, i.e. their ability to detect existing differences between subgraph probabilities. Previous studies have used the empirical FWER as a proxy for statistical power [1,2]. The rational underlying this approach is that the more conservative a method is (i.e. the more the actual FWER falls below the desired significance level), the lower its statistical power. In the following we will take a more direct approach and evaluate the performance of the methods under the alternative hypothesis of unequal subgraph probabilities. Again we will utilize Erdős-Rényi models, only now with different connection densities *p*_1_ = *p*_2_ for the two graph-generating processes. The question is: How many subgraphs are we able to correctly identify as being generated with distinct probabilities by the two processes? The answer to this question will not only depend on the multiple comparisons correction used but also on the sample size, the network size, and the effect size. The effect size *for a particular subgraph* G can be identified with the magnitude of the difference of subgraph probabilities 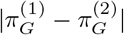. The larger this difference, the better the chances to detect the effect. In the following we will use the difference between the connection densities *p*_1_ and *p*_2_ as a measure of the effect size for the entire graph-generating processes.

In a simulation study we analyzed sample sizes n = 20, 30, 40. We set the probability of individual links for the first graph-generating process to *p*_1_ = 0.2. The second process generated individual links with probability p_2_ = 0.2 + e, where *e* = 0.1, 0.2, 0.3. Since *p*_1_ and p_2_ are chosen smaller than or equal to 0.5, the effect sizes for particular subgraphs are a decreasing function of the number of edges they consist of. In other words, the difference is more pronounced for subgraphs consisting only of few edges and can become very small for complex subgraphs. We considered network sizes *l* = 2, 4, 6, 8,10. For each possible choice of n, e, and l we simulated 1000 data sets and applied significant subgraph mining with either Tarone, Westfall-Young or Bonferroni correction. The number of permutations for the Westfall-Young procedure was set to 10000 as recommended in previous studies [2]. The two graph-generating processes were sampled independently (between subjects case) and accordingly Fisher’s exact test was utilized. The results are shown in Figure 6.

**Fig 6.**
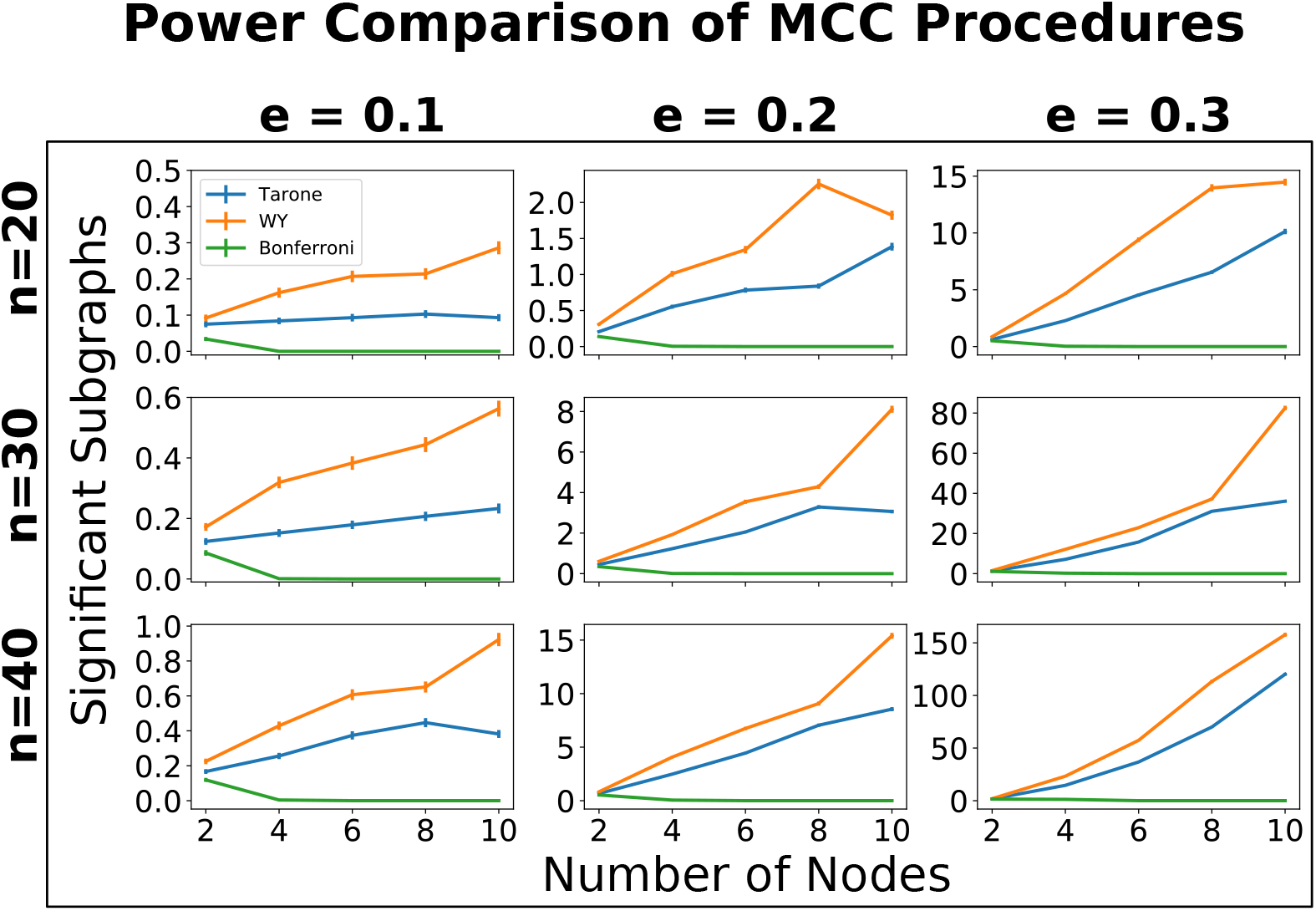
Average number of significant subgraphs identified depending on correction method, samples size, network size, and effect size. Error bars represent one standard error. The number of identified subgraphs increases with sample size (rows) and effect size (columns) for all correction methods.

As expected the average number of detected significant subgraphs is an increasing function of both sample size and effect size. The relationship between detected differences and number of nodes is less straightforward. Generally there is an increasing relationship, but there are a few exceptions. The likely explanation for this phenomenon is that there is a trade-off between two effects: on the one hand, the larger the number of nodes the more differences there are to be detected. But on the other hand, the larger the number of nodes the more severe the multiple comparisons problem becomes which will negatively affect statistical power. For some parameter settings this latter effect appears to be dominant. The most powerful method is always the Westfall-Young correction followed by the Tarone correction. The Bonferroni correction has the worst performance and its power quickly drops to zero because the corrected threshold can in principle not be attained.

Generally, only a very small fraction of existing differences is detectable. Since the graphs are generated by independently selecting possible links with a fixed probability, the subgraph probability is a decreasing function of the number of links a subgraph consists of. Complex subgraphs are quite unlikely to occur and will therefore not be testable. Additionally, the difference between subgraph probabilities 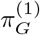 and 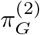 decreases with increasing subgraph complexity making this difference more difficult to detect. For instance, if e = 0.3, then the difference in subgraph probabilities for subgraphs with 10 nodes is about 0.001. Accordingly, even with a sample size of 40, only a small fraction of existing differences is detectable.

## Empirical Power Analysis with Transfer Entropy Networks

We applied the subgraph mining method to a data set of resting state MEG recordings comparing 20 autism spectrum disorder patients to 20 neurotypical controls. The details of the study are described in [12]. Here, seven voxels of interest were identified based on differences in local active information storage; subsequently timecourses of neural mass activity in these voxels were reconstructed by means of a linear constraint minimum variance (LCMV) beamformer. The locations of the voxels are shown in Table 1. We applied an iterative greedy method to identity transfer entropy networks on these voxels ( [13, 14]). The goal of this method is to find for each target voxel a set of source voxels such that 1) the total transfer entropy from the sources to the target is maximized, and 2) each source provides significant transfer entropy conditional on all other source voxels in the set. The outcome of this procedure is one directed graph per subject where each link represents significant information transfer from one voxel to another (conditional on the other sources). Accordingly, we are in a setting in which subgraph mining is applicable. The inferred transfer entropy graphs are shown in Figures 7, 8. Note that the edges are labeled by numbers that represent the time lags at which information transfer occurred. The parameters of the network inference algorithm were chosen so that lags are always multiples of five. Since the sampling rate was 1200Hz this corresponds to a lag increment of ≈ 4ms. So the graph representation also contains information about the temporal structure of information transfer and differences in this structure can be detected by subgraph mining as well. For example, even if the probability of detecting information transfer from voxel 0 to voxel 1 is the same in both groups, this transfer may be more likely to occur at a time lag of 5 (≈ 4ms) in the autism group whereas it may be more likely to occur at a time lag of 10 (≈ 8ms) in the control group.

**Fig 7.**
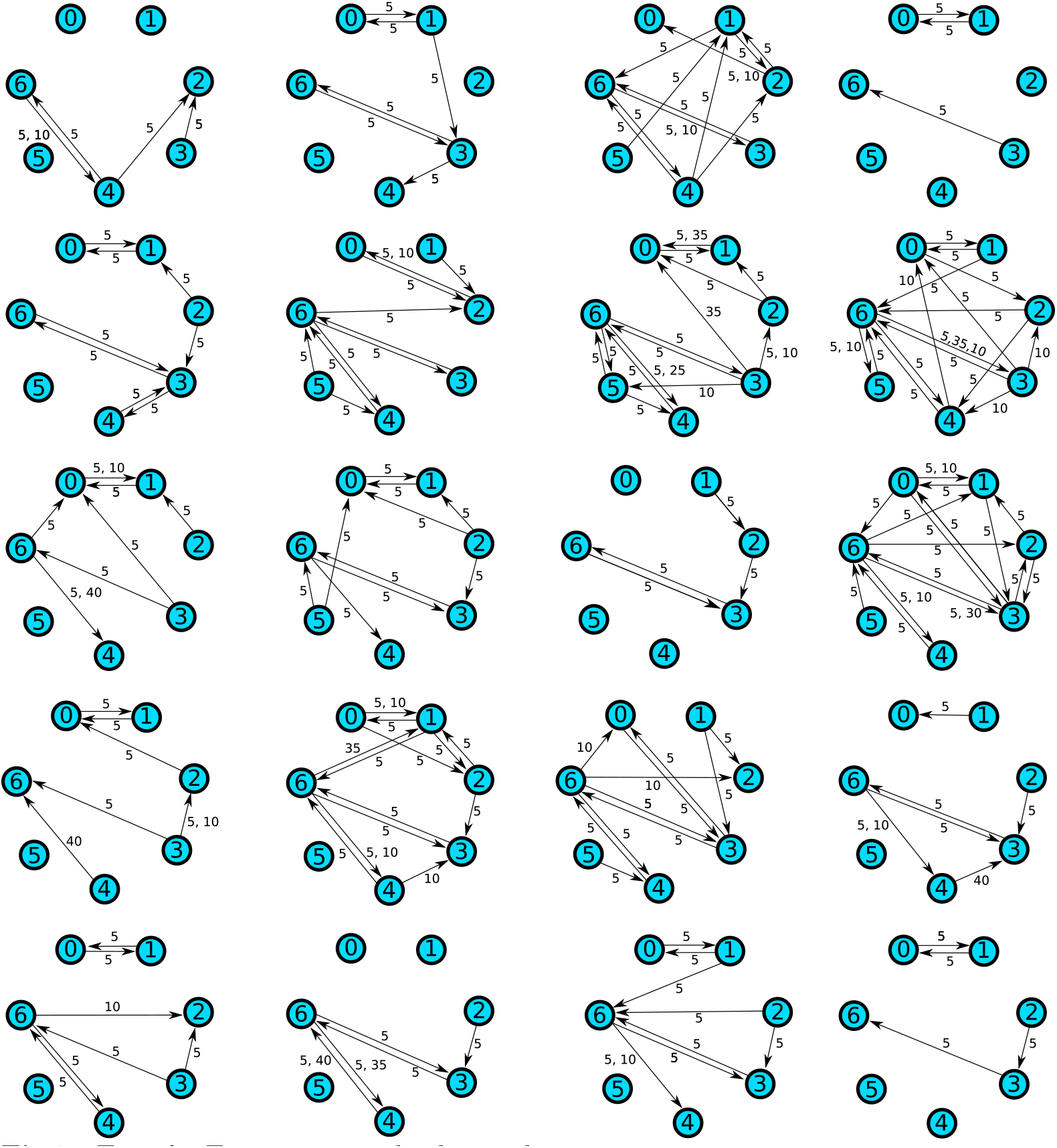
Transfer Entropy networks detected in autism spectrum group.

**Fig 8.**
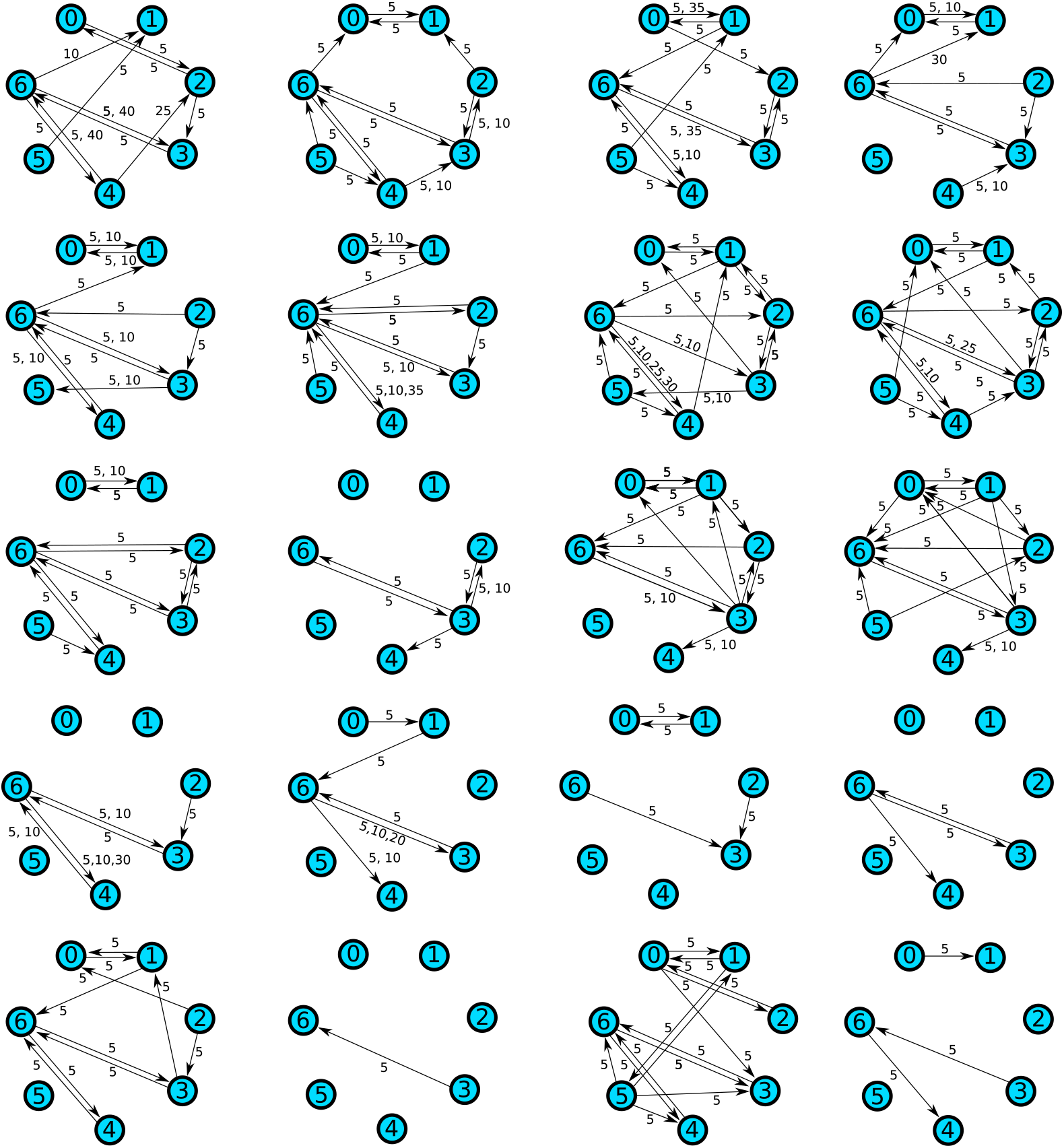
Transfer Entropy networks detected in control group.

**Table 1.** Voxel IDs and corresponding brain regions.

| Voxel ID | Location |
| --- | --- |
| 0 | Cerebellum |
| 1 | Cerebellum |
| 2 | Lingual Gyrus / Cerebellum |
| 3 | Posterior Cingulate Cortex (PCC) |
| 4 | Precuneus |
| 5 | Supramarginal Gyrus |
| 6 | Precuneus |

We applied subgraph mining with both Tarone and Westfall-Young correction to this data set. No significant differences between the ASD group and control group could be identified. Due to the rather small sample size, this result is not entirely unexpected. For this reason, we performed an empirical power analysis in order to obtain an estimate of how many subjects per group are required in order to detect existing differences between the groups. This estimate may serve as a useful guideline for future studies. The power analysis was performed in two ways: First, by resampling links independently using their empirical marginal frequencies, and second, by resampling from the empirical joint distribution. The latter can be achieved by first sampling a link with its empirical marginal probability. If the link occurs, then the second link is then sampled with its empirical conditional probability given that the first link occurred. If the first link does not occur, then the second link is then sampled with its empirical conditional probability given that the first link did not occur. This process is then repeated for the remaining links as well. For instance, given that the first link occurred and the second did not occur, the third link is sampled with the corresponding conditional probability and so forth. Note that this procedure is equivalent to randomly drawing networks from the original data sets with replacement.

The results of the power analysis assuming independent links are shown in Figure 9. We simulated sample sizes 20, 40, and 60 per group and carried out 400 simulations for each setting. The first notable outcome is that the original data are strikingly different from the results seen in independent sampling of links. In particular, the number of testable graphs is far higher in the original data (1272) than in the independently resampled data (28.7 on average and 55 at most). This indicates strongly that the processes generating the networks in ASD patients as well as controls do not generate links independently. Rather, there seem to be dependencies between the links such that some links tend to occur together making it more likely that subgraphs consisting of these links will reach testability. Accordingly, in the case of independent resampling much larger sample sizes are needed in order to detect the differences between the groups. Even in the *n* = 60 per group setting there were only 0.26 (Tarone) and 0.45 (Westfall-Young) significant subgraphs on average. There was no simulation in which more than three significant subgraphs were detected.

**Fig 9.**
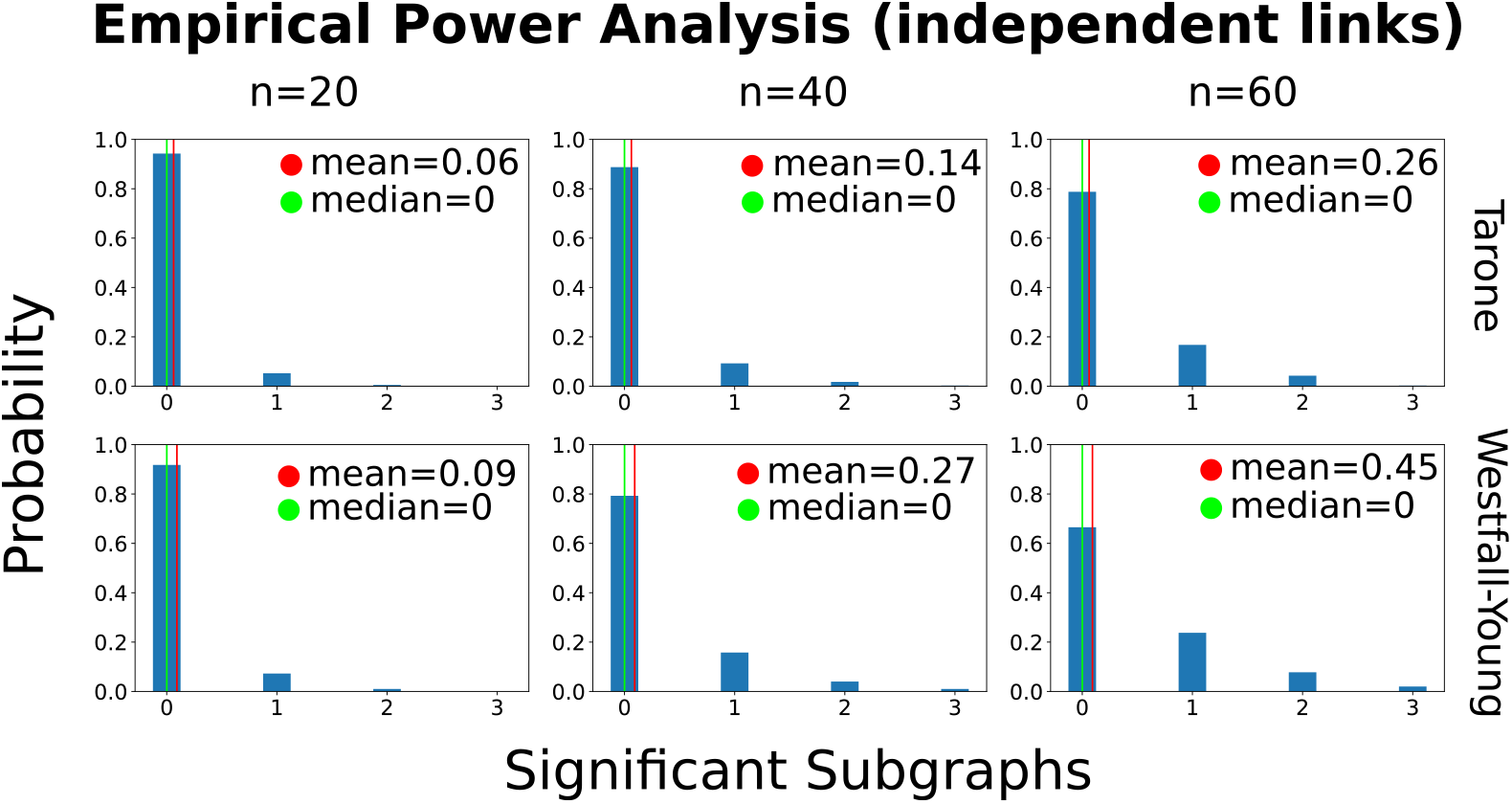
Results of empirical power analysis assuming *independence* of links. We simulated sample sizes 20, 40, and 60 per group and carried out 400 simulations in each setting. The histograms describe the fractions of simulations in which different numbers of significant subgraphs were detected.

The simulation results of the empirical power analysis based on the empirical joint distribution are shown in Figure 10. Again we used sample sizes 20, 40 and 60. The average number of testable subgraphs is in the same order of magnitude as in the original data set for the *n* = 20 setting (≈ 5200). Moreover, the number of identified significant subgraphs is far greater than in independent sampling for all sample sizes. The Westfall-Young correction identifies more subgraphs on average than the Tarone correction: 17.41 compared to 0.86 for *n* = 20, 202.20 compared to 14.88 for *n* = 40, and 831,24 compared to 100.62 for *n* = 60. The distributions are always highly skewed with more probability mass on smaller values. This is reflected in the median values also shown in the figure. For example, notwithstanding the average value of 14.88 significant subgraphs in the n = 40 setting with Tarone correction, the empirical probability of not fining any significant subgraph is still ≈ 42%. For the Westfall-Young correction this probability is only ≈ 1.8% in the n = 40 setting. In the *n* = 60 setting both methods have high empirical probability to detect significant differences. In fact, the Westfall-Young correction always found at least one difference and the Tarone correction only failed to find differences in 2.5% of simulations. The total number of detected differences can be in the thousands in this setting.

**Fig 10.**
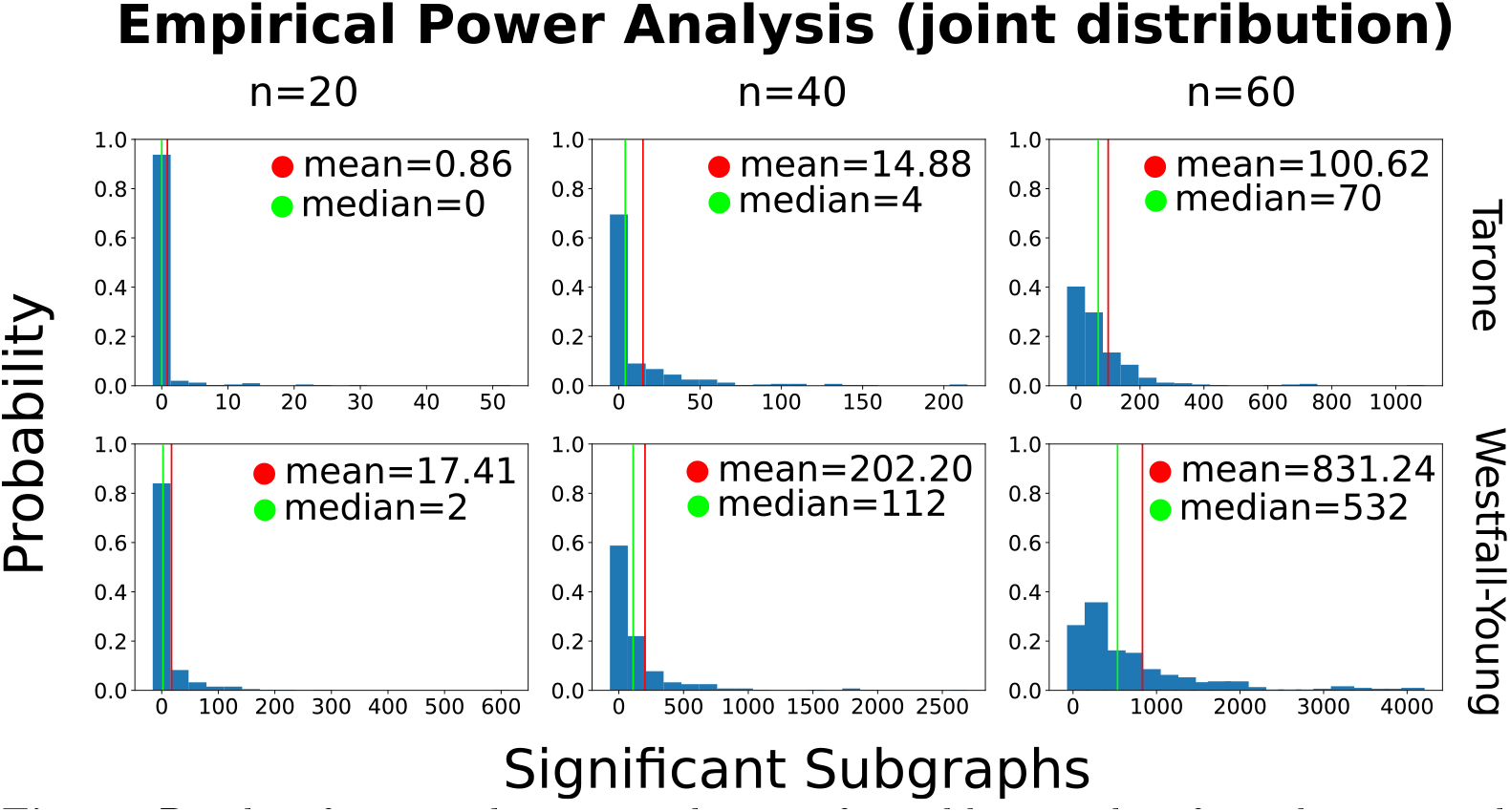
Results of empirical power analysis performed by sampling from the empirical joint distribution. We simulated sample sizes 20, 40, and 60 per group and carried out 400 simulations in each setting. The histograms describe the fractions of simulations in which different numbers of significant subgraphs were detected.

Since in the *n* = 60 setting both methods are likely to detect some of the existing difference we performed a subsequent analysis to narrow down the effect sizes that can be detected in this case. For each possible effect size (any multiple of 0.05 up to 0.35) we enumerated all subgraphs with this effect size and calculated their empirical detection probabilities among the 400 simulations. In total there were about 3.7 million graphs occurring with different empirical probabilities in the two groups. Most of these (99.5%) are graphs that occur exactly once in the entire data set. One important reason for this phenomenon is the following: suppose a network contains a subgraph that occurs only once in the data set. Then removing any other edges or combination of edges from the network will again result in a subgraph that only occurs once in the data set. Consider for example the last network in the second row in Figure 7. It contains a connection from node 6 to node 3 at a lag of 35 samples. This connection does not occur in any other network. This means that if any combination of the other 18 links occurring in the network is removed, the result will again be a uniquely occurring subgraph. There are 2^18^ = 262144 possibilities for doing so in this case alone.

The averages of the empirical detection probabilities for each effect size are shown in Figure 11 (upper plots). An interesting outcome is that the detection probability is not a strictly increasing function of the effect size. Rather there is a slight drop from effect sizes 0.25 to 0.3. Given the standard errors of the estimates this result might still be explained by statistical fluctuation (the two standard error intervals slightly overlap). However, in general this type of effect could also be real because the effect size is not the only factor determining detection probability. This is illustrated in Figure 11 (lower plots) which shows average detection probability over the smaller of the two occurrence probabilities 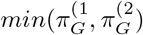. It turns out that the more extreme this probability is, the more likely the effect is to be detected. The highest detection probability is found if the empirical probability of occurrence is zero in one of the groups. For this reason it can in fact be true that the detection probability is on average higher for effect sizes of size 0.25 than 0.3, if the absolute occurrence probabilities are more extreme in the former case. In the data analysed here this is in fact the case: roughly half of the subgraphs with effect size 0.25 do have occurrence probability zero in one of the groups whereas this is not true for any of the subgraphs with effect size 0.3.

**Fig 11.**
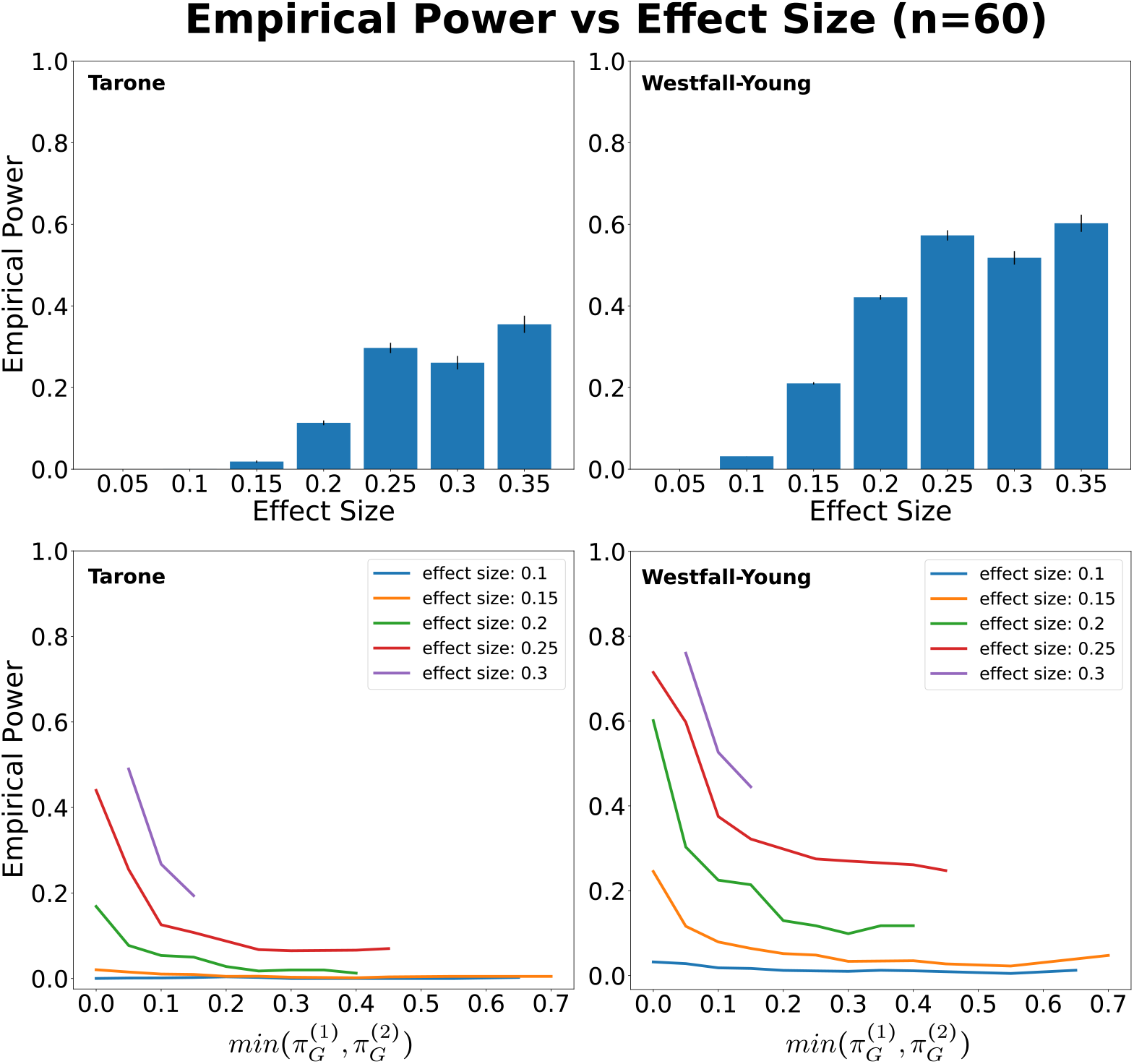
Upper plots: Average empirical detection probabilities for subgraphs with different effect sizes (i.e. the average is over all subgraphs with a certain effect size and for each particular graph the detection probability is estimated as the fraction of detection among the 400 simulations). Error bars are plus minus one standard error. Standard errors were not calculated for effect size 0.05 due to computational constraints. There are more than 3.7 million subgraphs with this effect size meaning that in the order of 10^12^ detection covariances would have to be computed This is necessary because the detections of different subgraphs are not independent. However, due to the this large number of subgraphs the standard errors are bound to be exceedingly small in this case. Lower plots: dependence of average detection probability on minimum of the two subgraph occurrence probabilities for different effect sizes. Even subgraphs with the same effect size have considerably different detection probabilities depending on how extreme the absolute occurrence probabilities are.

## Discussion

### What are the appropriate application cases for subgraph mining?

A key feature of significant subgraph mining that distinguishes it from other statistical methods for graph comparisons is that it considers all possible differences between graph generating processes. In other words, as soon as these processes differ in any way, subgraph mining is guaranteed to detect these differences if the sample size is large enough. This is in contrast to methods that only consider particular *summary statistics* of the graph generating processes such as the average degree of a node. Such methods are of course warranted if there is already a hypothesis about a specific summary statistic. For example, [15] were specifically interested in the entropy of the distribution of shortest paths from a given node to a randomly chosen second node. In such a case, performing a statistical test with respect to the statistic of interest is preferable over subgraph mining because the multiple comparisons problem is avoided. This leads to a higher statistical power *regarding the statistic in question*. On the other hand, the test will have a low power to detect other differences between the processes. Subgraph mining is the method of choice if no hypothesis about some specific aspect of the graph generating processes is available. It systematically explores the entire search space of all possible differences. This of course comes at the price of requiring a larger sample size.

### What are the requirements on sample size?

The appropriate sample size depends primarily on the kinds of effect sizes one seeks to be able to detect. Our empirical power analysis of the MEG data set discussed in the previous section suggests that in similar studies a sample size of about 60 is sufficient to have a very high probability to detect at least some of the existing differences. We carried out an additional analysis in order to narrow down the effect sizes likely to be detected at this sample size. This analysis showed that the largest effect sizes occurring in the empirical joint distribution (≈ 0.35 difference in probability of occurrence) had a detection probability of ≈ 0.4 on average using the Tarone correction and ≈ 0.6 on average using the Westfall-Young correction. This means that for a *particular* graph with a certain effect size the probability of detecting it is not extremely high. However, since there is generally a large number of such graphs there is a high probability of detecting at least some of them. Our analysis also showed that the effect size, understood as the difference in probability of occurrence of a subgraph between the groups, is not the only factor determining statistical power. Even graphs with the same effect size can have different probabilities of detection depending on how extreme the absolute probabilities of occurrence are. The detection probability is particularly high if the occurrence probability of a subgraph is close to zero in one of the groups. By symmetry we also expect this to be the case if it is close to one.

A possible way to reduce the amount of data required is to restrict the subgraph mining to subgraphs *up to a prespecified complexity*. For example, one could perform subgraph mining for all possible subgraphs consisting of up to three links. The validity of the method is not affected by this restriction. However, the search space is reduced and hence the multiple comparisons problem becomes less severe. In applying subgraph mining in this way it is important to pre-specify the desired complexity. Otherwise, we would run into yet another multiple comparisons problem. Consider the MEG data set presented in the previous section. Upon not detecting any differences with the full subgraph mining algorithm which considers all subgraphs on the seven nodes in our networks, one could check for differences among subgraphs consisting of at most six nodes. If nothing is found here either, we could move on to five nodes and so forth until we are down to a single link comparison. However, this approach would not be valid because the individual links are essentially given seven chances to become significant so that our bounds on the family-wise error rate do not hold anymore.

### What are the computational costs of subgraph mining?

Besides the required sample size another factor for the applicability of subgraph mining is the computation time. The number of possible subgraphs can very easily be large enough that it becomes impossible carry out a test for each one of them. Of course, the main idea behind the multiple comparisons methods presented here is that a large number of subgraphs can be ignored because they do not occur often enough or too often to be testable. For how many subgraphs this is true depends in particular on the connection density of the graphs. Generally, the computational load be will greater, the more nodes the graphs consist of and the more densely these nodes are connected. However, if the graphs are extremely densely connected one could revert to the *negative* versions of the graphs which would in this case be very loosely connected.

We provide a python implementation of significant subgraph mining as part of the IDTxl toolbox http://github.com/pwollstadt/IDTxl [16]. It offers both Tarone (with or without Hommel improvement) and Westfall-Young corrections. The latter is implemented utilizing the “Westfall-Young light” algorithm developed by [2] for between-subject designs. Details on the computational complexity can be found in this reference as well. The algorithm performs computations across permutations and achieves substantially better runtimes than a naive permutation-by-permutation approach. Our implementation is usable for both between-subjects and within-subject designs and allows the user to specify the desired complexity of graphs up to which subgraph mining is to be performed (see previous paragraph).

### Which multiple comparisons correction method should be used?

Regarding the choice between the two multiple comparisons correction methods we favor the Tarone correction because of its stronger error control guarantees. The Westfall-Young correction can at present only be assumed to provide weak control of the FWER. However, this makes it impossible to *localize* differences between graph generating processes. It only allows the conclusion that there must be *some* difference. The reason is essentially the same as the reason why it is not warranted to reject a null-hypotheses if is p-value has not been corrected for multiple comparisons at all. Suppose we perform 20 tests at level 0.05 and a particular null hypothesis, say the fifth one, turns out to reach significance. If we did not correct for multiple comparisons it would be a mistake to reject the fifth hypothesis because there is a plausible alternative explanation for why it reached significance: because we did not control for performing twenty tests, it was to be expected that we would see at least one hypothesis rejected and it just *happened* to be fifth one. Similarly, if we only have weak control of the FWER and a particular subgraph, say *G*_5_, reaches significance, then it would be a mistake to conclude that *G*_5_ is actually generated with difference probabilities by the two processes. The alternative explanation is that our false positive probabilities are not controlled under the actual scenario (the ground truth) and G5 simply happened to turn out significant. The only scenario that weak control *does* rule out (and this is how it differs from not controlling at all) is the one where all null-hypotheses are true, i.e. the one where the two graph generating processes are identical.

## Conclusion

Significant subgraph mining is a useful method for neural network comparison especially if the goal is to explore the entire range of possible differences between graph generating processes. The theoretical capability to detect any existing stochastic difference is what distinguishes subgraph mining from other network comparison tools. Based on our empirical power analysis of transfer entropy networks reconstructed from an MEG data set we suggest to use a sample size of at least 60 subjects per group in similar studies. The demand on sample size and computational resources can be reduced by carrying out subgraph mining only up to a prespecified subgraph complexity or by reverting to the negative versions of the networks under consideration. The method can also be used for dependent graph generating processes arising in within-subject designs when the individual hypothesis tests and multiple comparisons correction methods are appropriately adapted. We provide a full python implementation as part of the IDTxl toolbox that includes these functionalities.

## Acknowledgments

AG and MW are employed at the Campus Institute for Dynamics of Biological Networks (CIDBN) funded by the Volkswagen Stiftung. MW received support from the Volkswagenstiftung under the programme ‘Big Data in den Lebenswissenschaften’. This work was supported by a funding from the Ministry for Science and Education of Lower Saxony and the Volkswagen Foundation through the “Niedersachsisches Vorab”. MW received support from CRC 1193 C04 funded by the DFG. We thank Lionel Barnett for helpful discussions on the topic.

## Footnotes

1 Note that in the extreme case that the subgraph always occurs the resulting hypergeometric distribution has only one mass point at *f*_1_ (*G*) = *n*_1_. In this case both 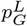 and 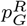 are equal to 1 and hence *p_G_* = 2. Similarly, if the subgraph does not occur at all, then there is only one mass point at *f*_1_(*G*) = 0. Again, *p_G_* = 2.

## Notes

### Competing Interest Statement

The authors have declared no competing interest.

